# LANA-dependent Interaction of Host Factors DAXX and BRD4 Impact KSHV Lytic Replication

**DOI:** 10.64898/2026.01.28.699085

**Authors:** Maria del Carmen Chacon Castro, Rithika Adavikolanu, Kelsey M. Haas, Yuan Zhou, Nevan J. Krogan, Robyn M. Kaake, Erica L. Sanchez

**Author notes:** Corresponding author: Erica L. Sanchez.

## Abstract

Kaposi’s Sarcoma-Associated Herpesvirus (KSHV) is an oncogenic gammaherpesvirus that causes Kaposi’s Sarcoma (KS). KSHV alternates between latent and lytic phases. Latency is marked by limited viral gene expression and absence of virion production, whereas the lytic phase is characterized by the expression of all viral genes in a temporally and sequentially regulated cascade of immediate-early, early, and late genes, culminating in viral genome replication and infectious virion production. Both KSHV latent and lytic viral phases contribute to its pathogenesis and the development of KS. KSHV proteins hijack host proteins, rewiring host cellular processes and signaling pathways, and modulating host gene expression. KSHV establishes latency through stable interactions between viral proteins and host factors, with the Latency Associated Nuclear Antigen (LANA) serving as a key regulator that interacts with host machinery to maintain the viral episome and regulate viral replication. However, the consequences for host proteins binding viral proteins, including changes in their interaction networks and their impact on viral replication, remains poorly understood. Here, using Immunoprecipitation Mass Spectrometry (IP-MS), we identified and validated the host death domain-associated protein (DAXX) and DNA ligase 3 (LIG3) as major LANA-associated factors and discovered a LANA-dependent recruitment of Bromodomain-containing protein 4 (BRD4) to DAXX. This remodeling of the DAXX interactome guided functional analyses demonstrating roles for DAXX and BRD4 in supporting KSHV infection. Functional studies show that siRNA knockdown of DAXX or BRD4 in iSLK.BAC16 cells, followed by lytic reactivation, significantly induced viral gene transcription and protein expression of KSHV lytic genes ORF45, ORF59, ORF26, and K8.1. Conversely, siRNA knockdown of LIG3 reduced the transcription and protein expression of KSHV lytic genes. Viral genome replication and infectious virion production were elevated upon knockdown of DAXX or BRD4 and reduced after the knockdown of LIG3. Additionally, chemical inhibition of BRD4 activity by the drug JQ1 in iSLK.BAC16 cells, followed by lytic reactivation, resulted in elevated KSHV lytic gene expression, genome replication, and infectious virus production. Together, these data suggest that both DAXX and BRD4 host genes contribute to KSHV latency maintenance, while LIG3 is required for lytic reactivation. Understanding the functional significance of these LANA-interacting host factors in regulating KSHV infection is critical for identifying therapeutic targets and developing potential treatment strategies.

## INTRODUCTION

Kaposi’s sarcoma-associated herpesvirus (KSHV), also known as human herpesvirus 8 (HHV8), is the etiological factor for Kaposi’s Sarcoma (KS), a common endothelial cell-based cancer in HIV/AIDS patients worldwide. KSHV is also associated with Primary Effusion Lymphoma (PEL), Multicentric Castleman’s Disease (MCD), and KSHV Inflammatory Cytokine Syndrome (KICS)^1^, making KSHV a significant global health concern. As a member of the gammaherpesvirus family, KSHV establishes lifelong infection in the host by alternating between two distinct phases, latency and lytic replication^2–4^. Latency is characterized by viral genome persistence as a circular episome, limited viral gene expression (5 genes), and absence of virion production, allowing the virus to persist in a dormant state within the host cell^5–8^. Periodic reactivation into the lytic cycle leads to widespread viral gene expression, genome replication, and the production of infectious virions, facilitating viral dissemination and contributing to pathogenesis^9–13^.

KSHV switches from latency to lytic cycles by exploiting host cellular machinery, such as manipulating host hypoxia responses, hijacking RNA processing mechanisms, and rewiring glutaminolysis, glycolysis, and fatty acid synthesis, to support viral gene expression, replication and survival^14–20^. Viral proteins, particularly those expressed during latency, interact with a diverse array of host factors to control gene expression, genome maintenance, immune evasion, and cell survival^21–23^. One of the highly-expressed latent proteins is the latency-associated nuclear antigen (LANA)^24^, which functions as a pleiotropic master regulator that maintains KSHV latency through the formation of LANA nuclear bodies by liquid-liquid phase separation. LANA tethers the viral episome to host chromatin by binding viral terminal repeats and interacting with host histones and chromatin proteins. Additionally, LANA recruits transcriptional silencers to repress lytic genes and assembles host replication machinery to ensure episome duplication and segregation^25–27^. The first AP-MS-derived KSHV-host protein-protein interaction (PPI) network from HEK293T cells characterized 556 stable host interacting proteins for 67 KSHV proteins, including 34 LANA interactors^28^. Among these LANA interactors were DAXX and LIG3, interactions that have been consistently reported. DAXX binding has been validated through imaging studies in overexpression systems showing DAXX recruitment to LANA nuclear bodies, and MS analyses and co-immunoprecipitation in both overexpression systems and infected PEL cells. Furthermore, LIG3 binding was validated in HEK293T cells through AP-MS analysis^28–31^. While the first AP-MS derived network did not identified BRD4 as a stable LANA interacting protein in HEK293T cells, targeted co-immunoprecipitation studies showed that FLAG-LANA pulled down endogenous BRD4 in C33A epithelial cells. Moreover, mapping studies in BJAB cells identified the ET domain within the C-terminal of BRD4 as the binding region for LANA. Furthermore, complementary NMR chemical-shift perturbation analysis using isotopically labeled LANA demonstrated that the BRD4 ET domain binds a single, defined site on the LANA C-terminal domain^32,33^. These studies primarily describe the interactions or their broader cellular consequences without addressing the specific functional contribution of these host factors to KSHV infection. DAXX plays essential roles in genome stability, transcriptional regulation, chromatin remodeling, apoptosis, and the DNA damage response, and its dysregulation has been linked to multiple cancer^34–38^. BRD4 is a bromodomain protein that regulates transcription, is associated with mitotic chromosomes, and has been implicated in cancer development^32,39–43^. LIG3 is a DNA ligase central to alternative non-homologous end joining, repairs DNA strand breaks, and defects in its activity contribute to genome instability and cancer progression^44–48^. Although these host factors each carry out distinct functions in chromatin regulation and DNA repair, their coordinated roles in KSHV infection remains unclear. While DAXX and BRD4 have both been shown to colocalize to LANA in the nucleus^29,32,49^, and LIG3 has been identified as a LANA-interacting protein^28^, the functional significance of these interactions in KSHV infection has not been defined. LIG3, in particular, has not been characterized in the context of KSHV replication. Despite evidence suggesting potential regulatory roles for these factors, how they influence viral gene expression, genome replication, virion production, or the balance between latency and lytic reactivation remains unclear.

Here, we show for the first time that BRD4 is recruited to DAXX in a LANA-dependent manner, revealing a previously unrecognized connection among these factors. No prior study has connected this interaction to the regulation of viral gene expression or virion production. In this study, we report how LIG3, DAXX and BRD4 contribute to KSHV infection by establishing their functional roles. Using the iSLK.BAC16 cell model, which harbors a recombinant KSHV genome and supports both latent infection and inducible lytic reactivation, we performed siRNA-mediated knockdown of these host genes followed by chemical induction of the lytic cycle. We found that depletion of DAXX or BRD4 resulted in enhanced early and late viral gene expression, increased viral genome replication, and elevated production and release of infectious virions, suggesting that these proteins act as host restriction factors that help maintain latency. Additionally, pharmacological inhibition of BRD4 reproduced the same effects seen with BRD4 knockdown, supporting its role in inhibiting lytic reactivation. Conversely, knockdown of LIG3 led to a significant reduction in lytic gene expression and virus production, implicating it as a positive regulator of lytic replication. Together, these findings demonstrate that LANA-interacting host proteins play distinct roles in regulating KSHV latency and lytic replication. By functionally characterizing the roles of DAXX, BRD4, and LIG3, our study provides new insights into the molecular interplay between KSHV and host factors and identifies potential therapeutic targets for the control of KSHV-associated diseases.

## MATERIAL AND METHODS

### Cells and Viruses

iSLK.BAC16 cells, stably maintaining a selectable GFP-expressing KSHV genome, and iSLK cells were maintained at 37°C and 5% CO_2_ atmosphere in iSLK media: Dulbecco’s modified Eagle medium (DMEM) (Corning, #10-013-CV) supplemented with 10% FBS (Corning, #35-011-CV) and 2 mM L-glutamine (Gibco, #25030-081). iSLK.BAC16 cells were selected with 1 µg/ml puromycin (Mirus, #MIR 5940), 0.1 mg/ml G418 (Fisher Bioreagents, #BP673-5), and 50 µg/ml hygromycin B (Corning, #30-240-CR) to achieve stable latent infection. iSLK.BAC16 cells were treated with 1 mM sodium butyrate (NaB) (Sigma Aldrich, #303410) and 1 μg/ml doxycycline hyclate (DOX) (Sigma Aldrich, #D5207) to induce lytic replication. For iSLK cells, iSLK media was supplemented with 1 μg/ml puromycin and 250 μg/ml G418. Experiments using iSLK.BAC16 and iSLK cells were performed using Dulbecco’s modified Eagle medium (DMEM) supplemented with 10% FBS and 2 mM L-glutamine. HEK293T cells (ATCC, #CRL-3216) were cultured and maintained in T175 flasks (Fisher, #12-556-011) at 37°C and 5% CO_2_ in DMEM with L-glutamine without sodium pyruvate (Fisher, #MT 10-017-CV) supplemented with 10% FBS (Life Technologies, #A3160502) and 1X Penicillin/Streptomycin (Fisher, #MT 30-002-CI).

### HEK293T Expression Constructs

LANA was subcloned from LANA-Strep-tagged pcDNA/TO (Invitrogen)^28^ into pQCXIP vector (Addgene) with a HTBH c-terminal tag. The LANA reading frame from pcDNA backbone was amplified and inserted into pQCXIP-HTBH using Gibson assembly. The LANA-HTBH plasmid was sequenced and matched against the reference sequence^28^. The empty pQCXIP-HTBH vector was a gift from the Huang lab^50^.

### Experimental Design and Statistical Rationale

For each of the four bait and control IPs (e.g. 1–DAXX, 2–LIG3, 3–mouse IgG, and 4–rabbit IgG), in both the LANA overexpression (LANA) and empty vector (EV) conditions, we performed 3 independent biological replicates (cells thawed, cultured, and transfected on different days) (**Table S1**). No technical or process duplicates were performed. In total, we collected 24 experimental samples for IP experiments: 12 EV and 12 LANA (from cells overexpressing LANA fused to a c-terminal 6xhistidine-TEVcleavage-biotin-6xhistidine tag). The number of controls were selected based on the total number of independent preparations for each primary antibody based on secondary antibody used (i.e. mouse or rabbit IgG), such that samples prepared and purified on separate days all include both mouse and rabbit control IPs. The mouse and rabbit IgG controls were combined and used for controls to score putative PPIs by SAINTexpress (v 3.6.3) to discriminate background from high confidence interactions^51^. DAXX and LIG3 baits and mouse and rabbit IgG controls in both +LANA and EV conditions were also analyzed by CompPASS^52^ for increased confidence in identifying PPIs from purification background.

### siRNA transfections

iSLK.BAC16 cells were seeded at 1x10^5^ cells/well into a 6-well plate. At 24 hours post-seeding, when cells reached 60-70% confluency, they were transfected with 25 pmol siRNA (siDAXX, GeneGlobe ID: GS1616; siBRD4, GeneGlobe ID: GS23476, or siLIG3, GeneGlobe ID: GS3980) (Qiagen, #1027416) or AllStars negative control siRNA (siASN, Qiagen, #1027295), one well per condition, using Lipofectamine RNAiMAX (Invitrogen, #13778-150) in Opti-MEM Medium (Gibco, #11058-021) following manufacturer’s instructions. Briefly, 30pmol siRNA was diluted in 150µl Opti-MEM medium and mixed with 9µl Lipofectamine RNAiMAX Reagent pre-diluted in 150µl Opti-MEM medium. After 5 minutes incubation at room temperature, the complexes were added to the cells in a final 250µl volume of siRNA-lipid complex per well. siRNA-transfected iSLK.BAC16 cells were incubated at 37°C and 5% CO_2_ atmosphere for 24 hours before inducing viral reactivation with 1μg/ml DOX and 1mM NaB.

### HEK293T cell culture, transfection, and cell harvest for IP-MS

HEK293T cells were seeded at 1.2x10^7^ cells in 15cm^2^ dishes. 24 hours post-seeding (∼60-70% confluency) cells were transfected with 10μg total DNA of the LANA and EV expression constructs (**Table S1**) with PolyJet In Vitro DNA Transfection Reagent (SignaGen Laboratories) at a 1:3 μg:μl ratio (plasmid:transfection reagent) according to the manufacturer’s recommendations. Briefly, plasmid mixtures and PolyJet reagent were separately prepared and diluted in 0.5 mL serum-free DMEM, then combined to 1 mL total and vortexed before incubating for 20 minutes at room temperature before adding dropwise to cells. Transfected cells were grown for 24 hours, then the media was removed, and the cells were dissociated from the 15-cm dishes using room temperature Dulbecco’s Phosphate Buffered Saline without calcium and magnesium (D-PBS) supplemented with 10 mM EDTA. Cells were pelleted by centrifugation at 400xg for 3 minutes and washed with 10 mL D-PBS and frozen at -80°C until needed for IP experiments.

### HEK293T cell lysis and immunoprecipitation (IP) and protein digestion

Frozen cell pellets were lysed on ice in 1 mL of freshly prepared cold native lysis buffers (IP buffer [50mM Tris-HCl, pH 7.5 at 4°C, 150mM NaCl, 1mM EDTA], supplemented with 0.5% Nonidet P 40 Substitute (NP40; Fluka Analytical), and cOmplete mini EDTA-free protease and PhosSTOP phosphatase inhibitor cocktails (Roche). To lyse cells, they were freeze-thawed a total of three times as follows: 1) first frozen cell pellets were gently resuspended in native lysis buffer at room temperature for 5-10 minutes until completely thawed and resuspended by pipette and then flash frozen; 2) in the second round lysates were thawed at room temperature for 7 minutes with shaking at 1100 RPM and then frozen a third time at -80°C; 3) samples were thawed a third time and then incubated rotating end-over-end for 30 minutes at 4°C. Post incubation, lysates were clarified by centrifugation at 13000 RPM at 4°C for 15 minutes, supernatants were collected in fresh 1.5 mL protein lo-bind tubes (Axygen) and cell debris discarded. A small amount of each lysate (50 μl) was reserved to determine protein abundance by Bradford assay. For each biological replicate, equal amounts of protein were aliquot into separate tubes and brought to 500 μl and mixed with the following amounts of antibody (**Table S1**): 1) either 1 μg of mouse (Invitrogen, #MA110407) or rabbit IgG (Proteintech, #30000-0-AP); 2) 2.5 μg α-DAXX (Invitrogen, #MA119731); or 3) 2.5 μg α-LIG3 (Proteintech, #26583-1-AP). Lysate-antibody mixture was incubated rotating end-over-end for 2 hours at 4°C. Pierce Protein A/G magnetic beads (50 μL slurry; Thermo Fisher, #88802) were washed four times in 1.0 mL wash buffer (IP buffer supplemented with 0.05% NP40) and then resuspended in 50 μL IP buffer. Lysate-antibody mixtures were mixed with freshly washed Protein A/G bead slurry and incubated rotating end-over-end for 2 hours at 4°C. After incubation the samples were put on a magnetic stand to collect the flow thru and to wash the beads two times with 1 mL wash buffer, and then two times with IP buffer. Protein-bound beads were then transferred to a clean tube and incubated in 50 μl of Reduction/Alkylation buffer (2M urea, 50mM Tris-HCl, pH 8.0 at room temperature, 1mM DTT, and 3mM Iodoacetamide) for 45 minutes in the dark rotating at 700 RPM at room temperature (ThermoMixer C). After incubation an additional 3mM DTT was added and incubated for 5 minutes at room temperature to quench the reaction. After quenching, 2 μg of sequencing grade trypsin (Promega) was added to the bead mixture and incubated overnight at 37°C in a water bath. Samples were placed on a magnetic tube rack to collect the peptide supernatant and peptides were acidified by adding 10% formic acid to a final concentration of 1%. Acidified peptides were desalted using C18 OMIX tips (Agilent Technologies, #A57003100) according to the manufacturer’s protocols. Briefly, samples were resuspended in 3% acetonitrile (ACN), 2% formic acid (FA). Tips were conditioned twice with 50% ACN, 0.1% FA and equilibrated twice with 0.1% FA. Samples were loaded by 10 aspirate-dispense cycles, washed twice with 0.1% FA, and eluted with 50% ACN, 0.1% FA followed by 90% ACN, 0.1% FA into a single tube. Desalted peptides were centrifuged in a speedvac to dry and stored at -80°C.

### Mass spectrometry data acquisition

Dried peptide samples were resuspended in 12μL 2% acetonitrile, 0.1% formic acid solution and analyzed by LC-MS/MS using an Easy-nLC 1200 (Thermo Fisher Scientific) interfaced via a nanoelectrospray source (Nanospray Flex) coupled to an Orbitrap Fusion Lumos Tribrid mass spectrometer (Thermo Fisher Scientific). Peptides were separated on a 150μm x 15cm 1.5μm PepSep column (Bruker) with a linear 120-minute gradient at a flow rate of 600nL/minute (details in **Table S2**). Buffer A consisted of 0.1% FA in water, and buffer B was 0.1% FA in 80% acetonitrile. For each cycle, one full MS scan (350-1350 m/z, at 240,000 resolution with a normalized AGC target (%) of 250) was followed by data-dependent MS/MS scans (data-dependent mode=cycle time; cycle time = 1 sec) (details in **Table S2**). Each immunoprecipitation sample was run once with a clean wash gradient run between each individual sample to reduce carry-over effects. For full description of all LC, MS acquisition, and tune parameters see **Table S2**.

### Peptide and protein identification and high confidence PPI scoring

All raw data files (.raw) were searched with MaxQuant (v 2.4.9.0)^53^ against the human proteome (Uniprot canonical protein sequences downloaded May 25, 2023) concatenated with a fully randomized decoy database, using a 0.01 peptide and protein false discovery rate. The following search parameters were used: 1) protease/enzyme was set to specific and Trypsin/P was selected with 2 max missed cleavages; 2) Carbamidomethyl (C) was as fixed modification; 3) Oxidation (M) and Acetyl (Protein N-term) as variable modifications with 5 max number of modifications (**Table S3**). For each bait in each condition, PPIs were determined by scoring with SAINTexpress (v 3.6.3)^51^ and filtered for high confidence using CompPASS^52^ with both mouse and rabbit EV and LANA samples being combined to use as controls (12 total). We applied a two step scoring strategy to determine the final list of reported interactors using two different scoring stringency cutoffs: 1) for each bait and condition, an identified protein must have a SAINTexpress Bayesian False Discovery Rate (BFDR) < 0.01; and 2) an identified protein was considered a high confidence interactor for that bait and condition, if it had a CompPASS wd percentile > 0.9 (for LIG3) or 0.95 (for DAXX). High confidence interactions for DAXX and LIG3 were separately mapped and visualized with cytoscape (v. 3.10.3)^54^.

### Differential interaction score (DIS) analysis of +LANA and EV conditions

To determine LANA- and EV-condition-specific interactions, we computed differential interaction scores (DIS) for all high confidence interactions determined for DAXX and LIG3 in either LANA or EV conditions (similarly as previously described^55–58^). The DIS is calculated as the difference between the EV and LANA interaction scores for a given bait.

*Bait-Prey DIS =(SaintScore.x + wd_percentile)_EV_ /2 - (SaintScore.x + wd_percentile)_LANA_ /2*

Here the interaction score is calculated for each bait-prey pair in each condition as the average of the SaintScore.x (SAINTexpress) and wd percentile (CompPASS). A DIS near 0 indicates an interaction that is confidently identified in both NT and +RNAse conditions, while a DIS near -1 or +1 indicates that interaction is more specific to LANA or EV conditions respectively.

### Reverse transcription quantitative PCR (RT-qPCR)

Knockdown efficiencies of targeting siRNAs were evaluated by RT-qPCR 60 hours after siRNA transfection. iSLK.BAC16 cells transfected with siRNAs were untreated or treated with 1 μg/ml DOX plus 1 mM NaB 24 hours post-transfection and harvested at 36 hours following lytic induction. Total RNA was extracted from cells following the PureLink™ RNA Mini Kit (Invitrogen™, # 12183025) protocol. Briefly, cells were washed with PBS and incubated for 5 minutes with 300µl of fresh lysis buffer containing 1% BME at 4°C. Cells were scraped from the cell culture plates and lysates were transferred to clean RNase-free microcentrifuge tubes. One volume 70% ethanol was added to each volume of cell homogenate and mixed. Mixtures were transferred to the spin cartridge and centrifuged at 12,000 g for 15 seconds at room temperature. Flow-through was discarded and the spin cartridge was loaded with 700µl wash buffer I and centrifuged at 12,000 g for 15 seconds at room temperature. After discarding the flow-through, the spin cartridge was washed with 500µl wash buffer II and centrifuged at 12,000 g for 15 seconds at room temperature twice. Finally, the spin cartridge was centrifuged at 12,000 g for 2 minutes at room temperature to dry the membrane and transferred to a clean RNase-free microcentrifuge tube. 40µl RNase-free water was added to the spin cartridge, incubated for 2 minutes and centrifuged at 12,000 g for 2 minutes at room temperature to elute the RNA. Samples were stored at -80°C until use. cDNA was synthesized from RNA samples according to the iScript™ Reverse Transcription Supermix (BIORAD, #1708841) protocol on a T100 Thermal Cycler BIORAD. Briefly, 1µg of total RNA was reverse-transcribed in a 20µl reaction system using 4µl of the 5X iScript™ Reverse Transcription Supermix and nuclease-free water to adjust the volume. The reaction conditions were 25°C for 5 minutes, 46°C for 20 minutes, and 95°C for 1 minute. The resulting cDNA products were stored at -80°C until use. Quantitative PCR of cDNA samples was carried out using PowerUp SYBR Green Master Mix (Applied Biosystems, #100029284) on a QuantStudio™ 3 Real-Time PCR System Applied Biosystems, using primers listed in Table S1. Briefly, each 20µl PCR reaction contained 10µl of 2X PowerUp SYBR Green Master Mix, 1µl forward and reverse primer (5µM each) (**Table S4)**, 1µl cDNA, and 8µl nuclease-free water. The cycling program was as follows: UDG activation for 2 minutes at 50°C, activation (Dual-lock DNA polymerase) for 2 minutes at 95°C, 40 cycles of denaturation for 15 seconds at 95°C and annealing/extension for 1 minute at 60 °C; followed by melt-curve analysis consisting of 15 seconds at 95°C (1.6°C/second), 1 minute at 60°C (1.6°C/second), and 15 seconds of dissociation at 95°C (0.15°C/second) to verify amplification specificity. Relative quantification of lytic mRNA expression was normalized to actin or tubulin and calculated using the delta-delta Ct method, comparing experimental conditions to control conditions (siASN-transfected iSLK.BAC16 cells after DOX and NaB treatment). Normalization was performed according to the following formulas: ΔCt = Ct _target gene_ – Ct _reference gene (actin or tubulin)_. ΔΔCt = ΔCt _experimental_ - ΔCt _control (siASN +DOX+NaB)_. Fold change was calculated as: FC = 2^-ΔΔCt^. Experiments included at least three independent biological replicates and qPCR analysis was performed using three technical replicates per sample. Statistical significance was assessed using one-way ANOVA.

### Quantification of intracellular KSHV genomic DNA by qPCR

iSLK.BAC16 cells were seeded at 1x10^5^ cells/well into a 6-well plate per condition, knocked down and reactivated by 1 μg/ml DOX and 1mM NaB treatment for 48 hours as described above. Cells were pelleted at 300 x g for 5 minutes at room temperature, supernatant was removed, and cells were analyzed immediately for quantification of the intracellular KSHV DNA copy number. DNA was extracted using the QIAamp DNA Blood Mini Kit (Qiagen, # 51104) after an RNAse A treatment as recommended by the protocol. Briefly, 20µl of protease was added to 200µl cells resuspended in PBS. Then, 4µl of RNAse A (100mg/ml) was added and mixed. Next, 200µl of buffer AL was added to the sample and vortexed by 15 seconds. Samples were incubated for 10 minutes at 56°C. Subsequently, 200µl of 100% ethanol was added to the sample and vortexed by 15 seconds. The mixture was applied to the QIAamp Mini spin column and centrifuged at 6 000 g for 1 minute. The flow through was discarded, and 500µl of buffer AW1 was added to the spin column and centrifuged at 6 000 g for 1 minute. After discarding the flow through, 500µl of buffer AW2 was added to the spin column and centrifuged at 20 000 g for 3 minutes. The spin column was placed in a clean microcentrifuge tube, and 200µl of buffer AE was added, incubated for 5 minutes, and centrifuged at 6 000 g for 1 minute. Extracted DNA samples were stored at -80°C until use. qPCR was performed using PowerUp SYBR Green Master Mix (Applied Biosystems, #100029284) on a QuantStudio™ 3 Real-Time PCR System Applied Biosystems, with LANA primers listed in **Table S4**. Linearized LANA-HTBH plasmid was used to generate a standard curve (10 – 10 x e^10 copies). After measuring plasmid DNA concentration, copy number calculations were performed according to the following formula: copies/µl = [X g/µl DNA plasmid concentration/(plasmid length in base pairs x 660)] x 6.022 x 10^23^. Double-stranded beta actin DNA standard (Origene, # HK200550), with known copies/µl was used to generate a standard curve (10 - 10 x e^7 copies). Briefly, each 10µl PCR reaction contained 5µl of 2X PowerUp SYBR Green Master Mix, 0.8µl forward and reverse primer (2.5µM each), 2µl cDNA, and 2.2µl nuclease-free water. The cycling conditions were as follows: Stage 1: 50°C for 2 minutes, 95°C for 10 minutes; Stage 2 (40 cycles): 95°C for 15 seconds, 60°C for 1 minute; and Stage 3 (dissociation curve): 95°C for 15 seconds, 60°C for 15 seconds, 95°C for 15 seconds. Standard curves (Ct vs. number of copies) were generated using the standards and used to calculate the copy number of each sample based on its Ct value. Viral intracellular copy number for LANA was normalized to Actin copy number usig the formula: Normalized LANA _copy number_ = LANA _copy number_ / Actin _copy number_. Data were presented as a fold change, FC = Normalized LANA _copy number (experimental)_ / Normalized LANA _copy number (control; siASN +DOX+NaB)_. Experiments were conducted with a minimum of three independent biological replicates, and qPCR analysis was performed in triplicates for each sample. Statistical analysis was carried out using one-way ANOVA.

### Production of KSHV and viral titer

iSLK.BAC16 cells were seeded at 1x10^5^ cells/well into a 6-well plate. After 24 hours, when cells reached 60-70% confluency, they were transfected with siASN, siDAXX, siBRD4, or siLIG3, one well per condition. At 24 hours post transfection, cells were treated with or without 1 μg/ml DOX and 1mM NaB for 48 hours to induce lytic reactivation. Supernatants from siRNA-transfected iSLK.BAC16 cells were collected and centrifuged at 300 × g for 10 minutes at 4°C. Cell-free supernatant containing the virus was collected and stored at −80°C until use. To titer the virions produced by the iSLK.BAC16 knockdown cells, 1x10^4^ iSLK cells were plated in a well/condition of a 12-well plate and then infected with 500µl of KSHV-containing, cell-free supernatants in the presence of 1 μg/ml Polybrene (EMD Millipore, #TR-1003-G). Spinoculation was carried out at 300 × g for 1 h at 37°C, then incubated for 4 hours at 37°C and 5% CO_2_ atmosphere with gentle rocking every 30 minutes. Following incubation, cells were washed with PBS (Corning, #21-040-CV) and the media was replaced with iSLK cells media. Cells were maintained at 37°C and 5% CO_2_ atmosphere for 48 hours, and infection was assessed by fluorescence microscopy (EVOS^TM^ FL color Imaging System) and flow cytometry based on GFP+ cells (see details below).

### Fluorescent microscopy

KSHV-infected iSLK cells were evaluated for GFP expression, as an indicator of viral infection. Fluorescent live images of iSLK cells after 48 hours of viral infection were adquired using the 10X objective on an EVOS™ FL color Imaging System (ThermoFischer). Each experimental condition was imaged six times per biological replicate. For the knockdown experiments, three independent biological replicates were performed, whereas four biological replicates were included for the JQ1 treatment experiments. Representative images from these replicates were presented. Number of GFP+ cells were quantified using ImageJ. The number of infected cells (GFP+) were normalized to the corresponding uninfected population (GFP-) for each condition. Data are presented as the average percentage of GFP+ cells for each condition relative to the siASN GFP+ control population. Statistical analysis was assessed using one-way ANOVA.

### Flow cytometry

iSLK cells were infected with supernantants from iSLK.BAC16 cells transfected with siDAXX, siBRD4, siLIG3, and siASN control, in the presence or absence of lytic reactivation. After 48 hours of infection, iSLK cells were washed with PBS, tripsinazed, and neutralized with iSLK media. Cells were centrifuged at 300g for 5 minutes at room temperature. Supernatants were removed, and cells were resuspended in 100µl FACS buffer (2% FBS in PBS). Cells were fixed by the addition of 120µl 4% PFA and incubated on ice for 15 minutes. Later, cells were washed with 2ml PBS and centrifuged at 300g for 5 minutes at 4°C. Supernatants were removed and cell pellets were resuspended in 500µl FACS buffer. Flow cytometry analysis was performed using the BD Fortessa SORP flow cytometer equipped with filters suitable for FITC detection (Excitation/Emission 498/517 nm). During data collection, a minimum of 10 000 events per sample were recorded to ensure statistical significance. Data were analized using FlowJo V.10.9. Cell populations were identified by gating on forwad scatter (FSC) and side scatter (SSC) to exclude debris and doublets. Quadrant gating differentiated GFP- and GFP+ cell populations. FITC- cells (Quadrant 1 and 4) corresponded to the GFP- cells, uninfected cell population, whereas FITC+ cells (Quadrant 2 and 3) corresponded to the GFP+ cells, infected cell population. For each condition, the percentage of infected cells was normalized to the corresponding uninfected cell population. Data are presented as the fold change in GFP+ cells for each condition relative to the siASN GFP+ control population. Data represent the average of eight biological replicates, and statistical analysis was assessed using one-way ANOVA.

### Immunofluorescence

iSLK.BAC16 cells were transfected with siDAXX, siBRD4, siLIG3, and the control siASN for 24 hours exactly as described above. Cells were treated with or without 1 μg/ml DOX and 1mM NaB to induce lytic reactivation for 48 hours. Cells were washed with PBS and fixed with 4% PFA for 10 minutes, followed by a PBS wash. Cells were permeabilized and blocked with 1% BSA 0.2% TritonX-100 in PBS for 10 minutes. Primary antibodies were incubated for 45 minutes at room temperature, followed by a PBS wash and secondary antibody incubation for 45 minutes protected from light. Primary antibodies: OF59 (Millipore, #MABF2752, 1:1000) and K8.1 (Millipore, #MABF2678, 1:1000). Secondary antibodies: Goat anti-Mouse IgG2b Cross-Adsorbed Secondary Antibody, Alexa Fluor™ 568 (Invitrogen, #A21144, 1:1000) and Goat anti-Mouse IgG1 Cross-Adsorbed Secondary Antibody, Alexa Fluor™ 647 (Invitrogen, #A21240, 1:1000). Cells were washed with PBS and stained with DAPI for 5 minutes followed by a PBS wash. Images were obtained by the Evident Scientific FV4000 Confocal Laser Scanning Microscope with a 20Xobjective, 0.80 numerical aperture, and 2.19X zoom magnification. DAPI was excited at 405 nm using 0.8% laser power, and emission was collected within a spectral detection range of 430–470 nm. FITC was excited at 488 nm with 0.3% laser power, and emission was collected between 500 and 540 nm. Alexa Fluor 568 was excited at 561 nm using 0.1% laser power, with emission detected from 580 to 630 nm. Alexa Fluor 647 was excited at 640 nm with 1% laser power, and emission was collected within a spectral detection range of 652–713 nm. Raw 16-bit images were processed by CellSens. Brightnes and contrast were adjusted for each chanel across conditions, and channels were merged.

### Western blot

iSLK.BAC16 cells transfected with siDAXX, siBRD4, siLIG3, and the control siASN for 24 hours and treated with or without 1 μg/ml DOX and 1mM NaB to induce lytic reactivation for 48 hours were lysed in Pierce RIPA buffer (Thermo scientific, #89900). Total protein levels for lysates were quantified using Pierce BCA Protein Assay Kit (Thermo Scientific, #23225) following the manufacturer’s specifications. For each knockdown-condition 30µg of total protein lysate was loaded onto 4-20% 10-well Mini-PROTEAN TGX Gel (BIORAD, #4561094) and transferred onto PVDF membranes (BIORAD, #1620264). The membranes were blocked for 1 hour on a rocker with 5% skim milk in a 0.1% TBST solution (1× Tris-buffered saline with 0.1% Tween 20) (Thermo scientific, #28360) and probed with primary antibodies diluted in 5% skim milk-TBST by rocking at 4°C overnight. Secondary antibodies were diluted in 5% skim milk-TBST and reacted on a rocker at room temperature for 2 hours. Immunolabeled proteins were visualized using Clarity Western ECL substrate and an Amersham ImageQuant™ 800 Western blot imaging system (Cytiva). Immunodetection was performed using mouse antibodies against ORF45 (Sigma Aldrich, #SAB5300153, 1:1000), K8.1 (Millipore, #MABF2678, 1:500), and rabbit antibodies against alpha tubulin (Proteintech, #11224-1-AP, 1:2000). Secondary antibodies included HRP conjugated goat anti-rabbit (Jackson Immuno Research Laboratory, #111-035-003, 1:20,000) or anti-mouse IgG (Jackson Immuno Research Laboratory, #115-035-003, 1:20,000). Experiments were performed in at least triplicate, and representative Western blot images are included.

### Drug inhibition

iSLK.BAC16 cells were seeded at 1x10^5^ cells/well into a 6-well plate, one well per condition. At 24 hours post-seeding, cells were treated with 100nM JQ1 (Selleckchem, #S7110), a BRD4 inhibitor, for 72 hours. At 24 hours post JQ1 (tert-butyl 2- [(9S)7 (4-chlorophenyl) -4,5,13-trimethyl-3 -thia-1,8,11,12 -tetrazatricyclo [8.3.0.02,6] trideca-2 (6),4,7,10,12-pentaen-9-yl] acetate) treatment, cells were reactivated with 1 μg/ml DOX and 1mM NaB for 48 hours in the presence or absence of 100nM JQ1. RNA was extracted using the PureLink™ RNA Mini Kit (Invitrogen™, # 12183025), and cDNA was synthesized according to the iScript™ Reverse Transcription Supermix (BIORAD, #1708841), as detailed above. qPCR was performed to determine the changes in viral relative mRNA expression using primers listed in **Table S4**. Cell-free supernatants were used for viral titer on iSLK cells as detailed above.

### Statistical Analysis

Figures and statistical analyses were generated using GraphPad Prism 10 software. Experiments were performed at least in triplicate. Data are presented as the mean ± SEM. Experimental conditions were compared to control conditions using ANOVA. Significant p-values are noted in each figure (* ≤ 0.05; ** ≤ 0.01; *** ≤ 0.001), ns denotes non-significant change.

## RESULTS

### Comparative PPI network for DAXX and LIG3 in +/- KSHV LANA conditions identifies BRD4 as LANA-dependent DAXX PPI

As known KSHV LANA interactors, DAXX and LIG3 may play a key role in regulating KSHV latency or reactivation, yet it is unclear if host DAXX and LIG3 protein-protein interaction (PPI) networks are rewired in the presence of LANA. To characterize the LANA-dependent changes in host protein interaction networks for DAXX and LIG3, we performed comparative endogenous pull-downs of DAXX and LIG3 from LANA-expressing and non-expressing HEK293T cells (Fig. 1A). To this end, we transiently transfected HEK293T cells with EV or C-terminally HTBH tagged LANA. DAXX and LIG3 proteins were individually purified by anti-DAXX and anti-LIG3 immunoprecipitation respectively in biological triplicate from EV and LANA HEK293T cell lysates, with IgG antibodies used for background controls. Purified samples were digested with trypsin, and the resulting peptides were analyzed by LC-MS/MS. Data was searched using MaxQuant^53^ (**Tables S3 and S5**).

**Figure 1.**
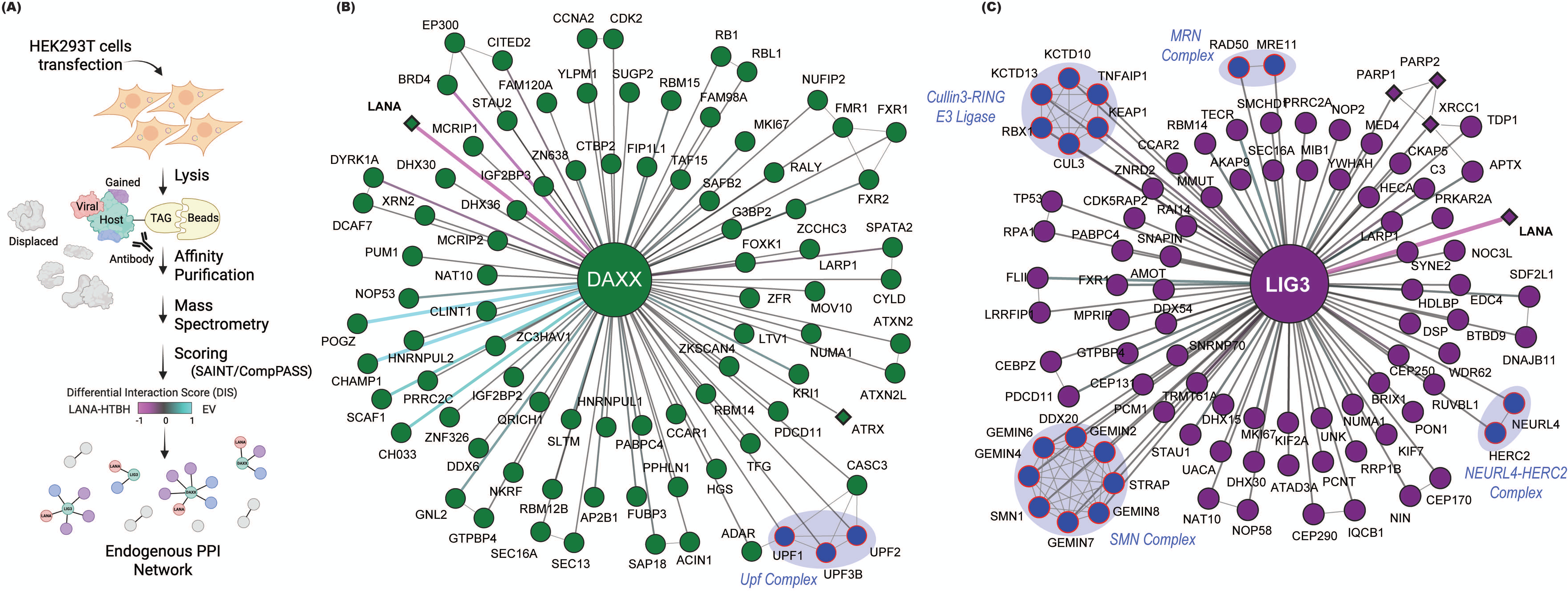
Comparative PPI network determination for DAXX and LIG3 in +/- KSHV LANA conditions. **A**. Schematic overview of the IP-MS and PPI determination strategy. Endogenous DAXX and LIG3 immunoprecipitation mass spectrometry (IP-MS) is carried out in +/- KSHV LANA conditions in triplicate. Proteins digested and peptides separated and analyzed by LC-MS/MS. MaxQuant identified peptides and proteins are scored to determine background from high confidence interactions using a two-step scoring approach. Step 1-proteins with SAINTexpress BFDR < 0.01; Step 2-proteins with CompPASS wd percentile > 0.9. Interactions are considered LANA-dependent if they have a DIS < -0.5 or > 0.5. **B-C**. Differential PPI networks for **B.** DAXX and **C.** LIG3. Circular nodes are high confidence human proteins that interact with DAXX or LIG3. Diamond nodes with thick borders have been reported in StringDB (https://string-db.org/)^95^ as physical interactors of DAXX or LIG3 with high confidence (0.9). Edge color indicates DIS from -1 (pink) to 1 (cyan). Grey edges are known interactions between the high confidence PPIs as reported in StringDB (https://string-db.org/)^95^ as physical interactors with high confidence (0.9). Nodes with red borders inside blue backgrounds are known complexes (labeled with blue italics text).

High-confidence PPIs were differentiated from background using a two-step filtering approach with the SAINTexpress^51^ and CompPASS^52^ scoring algorithms (**Fig 1A**). Proteins identified in each bait-condition were considered high confidence if they had a SAINTexpress BFDR <0.01. For LIG3 pull-downs, we used a second filter where the bait-prey CompPASS wd percentile score was > 0.9 (**Table S6**). DAXX pull-downs were somewhat higher in background, therefore we used a more stringent threshold of CompPASS wd percentile >0.95 as a filter (**Table S6**). In total, we identified 88 DAXX (82-EV; 58-LANA) and 94 LIG3 (85-EV; 76-LANA) high confidence interactors (Fig. 1B**-C**). Several proteins (13 shared PPIs) were identified as high confidence interactors in both DAXX and LIG3 pull-downs, likely because DAXX and LIG3 are both known to interact with chromatin and may share interactions as a result (**Table S6**). Notably, in our results, both DAXX and LIG3 pull down well-characterized interactors. For DAXX this included ATRX, and for LIG3 this included XRCC1, PARP1, and PARP2^59–72^.

While the majority of PPIs were identified in both conditions, some were identified as high confidence only under one condition. In some cases, these condition-specific interactions were pulled down and identified in both conditions, sometimes just below our stringent high confidence threshold. Therefore, to identify PPIs that are enriched in or specific to +/- LANA conditions, we used a differential interaction score (DIS)^55–58^. In this case, a DIS near 0 indicates that a specific DAXX or LIG3 PPI was confidently purified under both +/- LANA conditions, whereas a DIS of -1 or 1 indicates that a PPI is more specific to +LANA or EV conditions, respectively (**Table S6**). Using a -0.5<DIS<0.5 threshold, we did not identify any LANA-specific PPIs for LIG3, aside from LANA. For DAXX, however, we did identify 4 proteins that are EV specific (SFR19 or SCAF1, CH033, POGZ, CHAP1 or CHAMP1) and 1 protein (BRD4) which is recruited in the presence of LANA (Figure 1B**, Table S6**).

### DAXX and BRD4 knockdown enhance, while LIG3 knockdown suppresses KSHV lytic gene expression

To investigate the involvement of host factors in KSHV viral gene expression, we utilized iSLK.BAC16 cells, a well-established cell line that stably carries the BAC16 KSHV genome and expresses the viral lytic switch protein RTA (Replication and Transcriptional Activator) under the control of a doxycycline-inducible promoter, and encodes a constitutively expressed GFP reporter^73^. iSLK.BAC16 cells were transfected with siRNAs targeting DAXX and LIG3, host factors previously shown to interact with the latent viral protein LANA^28,29,49^, and BRD4, which was identified in our DAXX immunoprecipitation mass spectrometry analysis as interacting with DAXX only in the presence of LANA. Twenty-four hours post siRNA transfection, iSLK.BAC16 cells were treated with or without DOX and NaB to induce lytic reactivation. RNA was extracted after 36 hours of reactivation, and RT-qPCRs were performed to analyze the expression levels of viral genes. siRNA transfection with siDAXX, siBRD4, and siLIG3 in iSLK.BAC16 cells significantly reduced transcript levels by over 70%, compared to samples transfected with AllStars Negative control siRNA (siASN) (Figure 2A-C). Upon KSHV reactivation, lytic viral gene expression is a temporal and sequential process involving the expression of immediate-early, early, and late lytic genes^74^. To assess the impact of the knockdown of host factors DAXX, BRD4, and LIG3 on KSHV lytic gene expression, we measured the transcript levels of a representative immediate-early lytic gene ORF45, an early lytic gene ORF59, and late lytic genes ORF26 and K8.1 after reactivation upon siRNA treatment. After 36 hours of reactivation, the expression of the immediate-early lytic gene ORF45 was significantly increased in siDAXX- and siBRD4-treated cells, showing an 8-fold and a 14-fold increase, respectively, compared to the siASN control cells. In contrast, siLIG3-treated cells showed a trend toward reduced ORF45 expression compared to the siASN control cells (Figure 2D). Consistently, early lytic gene ORF59 expression was also significantly upregulated by 3-fold and 2.5-fold in siDAXX- and siBRD4-treated cells, respectively, compared to the siASN control cells, whereas siLIG3-treated cells showed a trending decrease of ORF59 expression compared to the siASN control cells (Figure 2E). Similarly, relative mRNA expression of the late lytic gene ORF26 was significantly elevated, exhibiting a 10-fold and 12-fold increase in siDAXX- and siBRD4-treated cells, respectively, compared to the siASN control cells (Figure 2F). Notably, relative mRNA expression of the late lytic KSHV gene K8.1 was significantly upregulated, with 20-fold and 37-fold increases in siDAXX- and siBRD4-treated cells, respectively, compared to the siASN control cells (Figure 2G). In contrast, siLIG3-treated cells exhibited downward trends in both ORF26 and K8.1 transcript levels compared to the siASN control cells (Figure 2F-G). While some leaky expression of lytic genes was detected during latency prior to reactivation, their levels were not statistically significant. Together, these results indicate that a loss of DAXX or BRD4 induces a significant increase in immediate-early, early, and late KSHV relative mRNA expression, suggesting that DAXX and BRD4 regulate latency maintenance. Conversely, depletion of LIG3 results in a consistent reduction in immediate-early, early, and late lytic KSHV genes relative mRNA expression, suggesting that LIG3 is required for efficient lytic reactivation.

**Figure 2.**
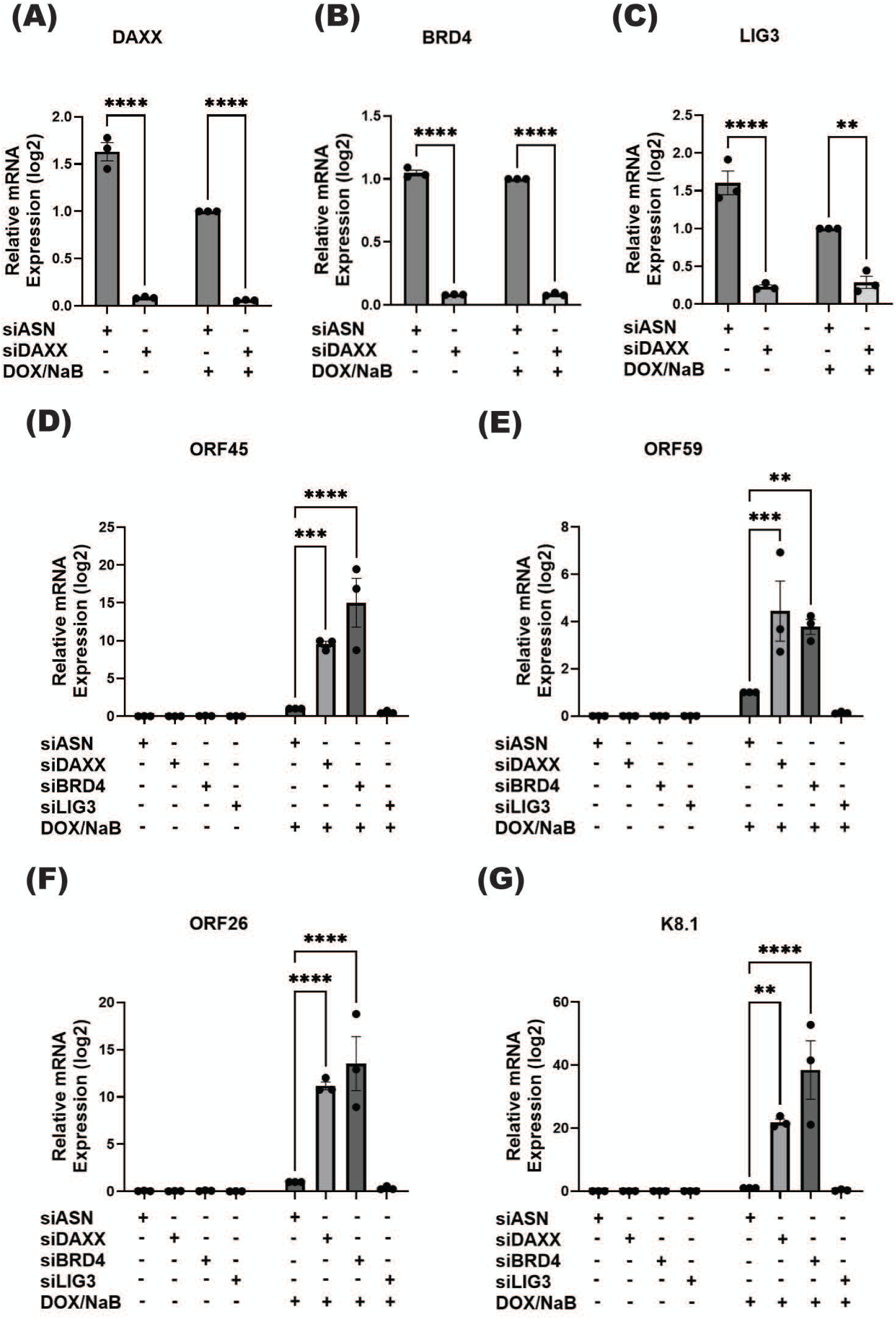
Knockdown of the host genes DAXX and BRD4 promotes increased, while knockdown of the host gene LIG3 promotes decreased KSHV lytic gene expression. Relative mRNA expression of the host genes (A) DAXX, (B) BRD4, (C) LIG3, and KSHV lytic genes (D) ORF45 (immediate early), (E) ORF59 (early lytic), (F) ORF26, and (G) K8.1 (late lytic) after the knockdown of the host gene DAXX, BRD4, and siLIG3 in iSLK.BAC16 cells treated with or without DOX/NaB. All expression data are relative to Actin or Tubulin (housekeeping) and normalized to siASN +DOX/NaB. siASN: AllStars negative control siRNA. ****, P≤0.0001; ***, P≤0.001; **, P≤0.01; *, P≤0.05; n=3, Error bars=SEM.

### DAXX and BRD4 inhibition increase while LIG3 inhibition decreases KSHV lytic protein expression

To confirm that the changes of KSHV lytic genes observed at the transcript level was also reflected at the protein level following the knockdown of the host factors DAXX, BRD4, and LIG3, we performed immunofluorescence analyses in iSLK.BAC16 cells after the siRNA treatments and 48 hours of lytic reactivation. KSHV lytic ORF45 and K8.1 protein levels were increased after DAXX and BRD4 inhibition compared to the siASN control samples, as observed in the representative confocal microscopy images. In contrast, LIG3 knockdown resulted in decreased protein expression levels of the KSHV lytic proteins ORF45 and K8.1 compared to the siASN control samples (Figure 3A). We also performed Western blot analyses after the knockdown of the host factors DAXX, BRD4, and LIG3 with or without lytic reactivation. These analyses revealed increased expression of lytic KSHV proteins ORF45 and K8.1 in siDAXX- and siBRD4-treated iSLK.BAC16 cells after lytic reactivation compared to siASN control cells, consistent with enhanced lytic reactivation. On the other hand, upon reactivation, siLIG3-treated cells showed decreased expression of lytic KSHV proteins ORF45 and K8.1 compared to siASN control cells (Figure 3B), consistent with impaired lytic reactivation.

**Figure 3.**
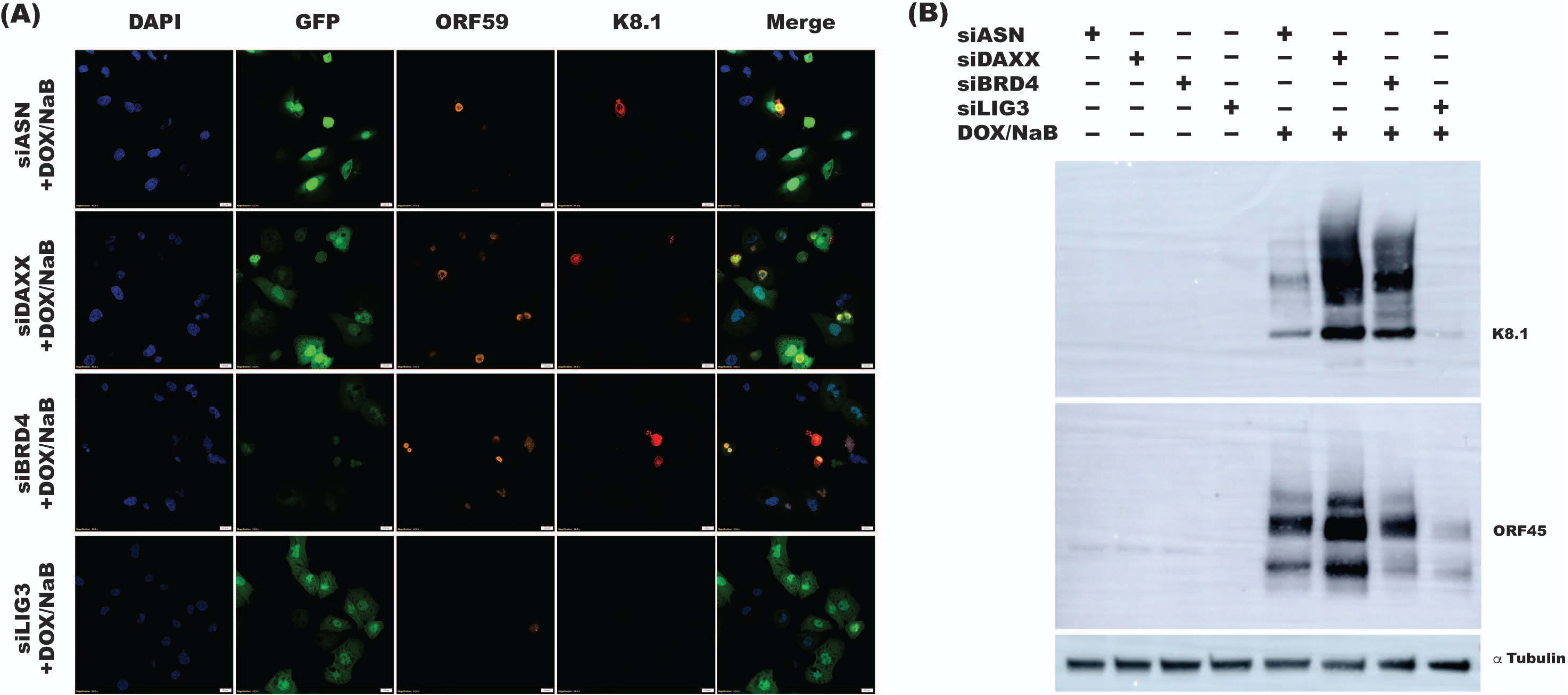
Knockdown of the host genes DAXX and BRD4 promotes increased, while LIG3 knockdown suppresses KSHV lytic protein expression. DAXX, BRD4, and LIG3 host genes were knocked down in iSLK.BAC16 cells followed by lytic reactivation were imaged by laser confocal microscopy. (A) Immunofluorescence staining of ORF59 (orange), K8.1 (red), GFP (expressed constitutively by iSLK.BAC16 cells) (green), and DAPI (blue) for the nucleus. (B) Protein levels of KSHV lytic genes were determined by Western blot. Representative Western blot pictures of early lytic protein ORF45 and late lytic protein K8.1 were included. Alpha Tubulin was used as the internal control. siASN: AllStars negative control siRNA.

### DAXX and BRD4 inhibition enhance while LIG3 knockdown suppresses KSHV genome replication

Viral KSHV DNA replication takes place before the transcription of late lytic genes^75,76^ (Figure 4A). To investigate the role of host factors DAXX, BRD4, and LIG3 in KSHV genome replication, we knocked down each host factor in iSLK.BAC16 cells using siDAXX, siBRD4, and siLIG3 and subsequently quantified KSHV LANA expression before and after lytic reactivation. We measured intracellular viral copy number by qPCR to assess the efficiency of viral genome replication in host cells. Viral copy number was normalized to actin copy number. Upon reactivation, intracellular viral copy number was significantly increased by 40% and 70% after the knockdown of DAXX and BRD4, respectively, compared to the siASN control. In contrast, our qPCR analyses show a trending decrease in intracellular viral copy number in the LIG3 knockdown iSLK.BAC16 cells after lytic reactivation compared to the siASN control (Figure 4B). Collectively, inhibition of DAXX and BRD4 gene expression by siRNAs followed by lytic reactivation resulted in an elevated viral copy number, which indicates an active and enhanced KSHV genome replication. Conversely, decrease in intracellular viral copy number after LIG3 knockdown in reactivated iSLK.BAC16 cells, suggest there is limited viral genome replication. Additionally, viral copy number before reactivation was consistent and low across all conditions, reflecting the latent phase during which episomal copy numbers are stably maintained.

**Figure 4.**
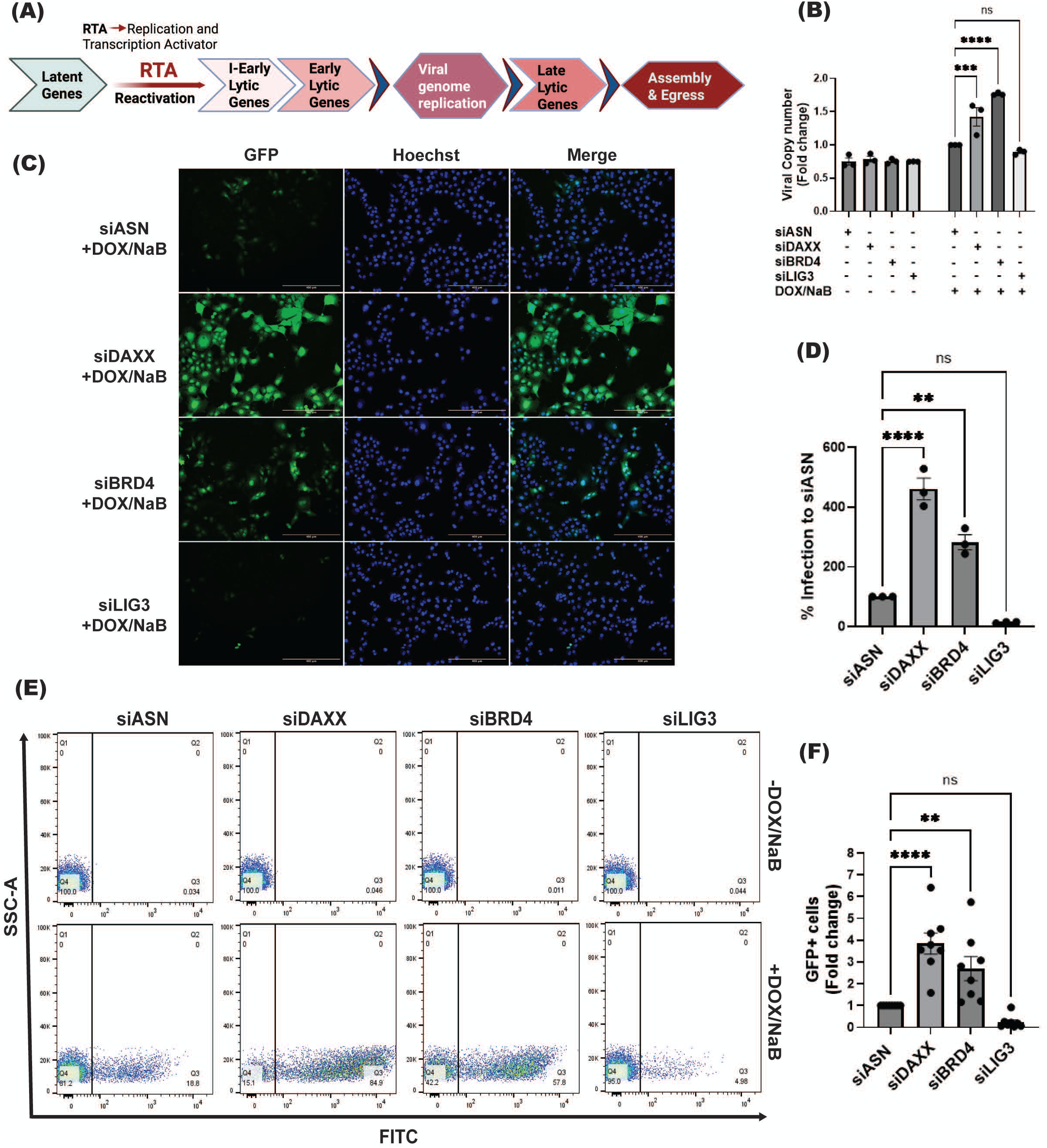
Host genes regulate KSHV genome replication and maximal KSHV virion production. (A) Schematic representation of the temporal and sequential KSHV replication cycle. Viral genome replication is an intermediate process after early lytic gene expression and before late lytic gene expression. (B) Intracellular viral copy number was calculated upon host gene knockdown by siDAXX, siLIG3, and siBRD4 in iSLK.BAC16 cells before and after lytic reactivation with DOX/NaB, n=3. Supernatants were collected after 48 hours of reactivation and viral titers were determined in iSLK cells on an EVOS fluorescent microscope at 10X (C). (D) Quantification of GFP + cells in (C), n=3. (E) Supernatants collected after host genes knockdown and 48 hours of reactivation of iSLK.BAC16 cells were titered on iSLK cells and also analyzed by Flow cytometry. (F) Quantification of GFP+ cells from (F), n=8. siASN: AllStars negative control siRNA. ****, P≤0.0001; ***, P≤0.001; **, P≤0.01; *, P≤0.05, Error bars=SEM.

### DAXX and BRD4 inhibition contribute to while LIG3 knockdown inhibits maximal KSHV virion production

Following the replication and packaging of the KSHV genome, fully assembled virions egress from the host cell, leading to the production of infectious virus particles during the final step of the lytic cycle^77–80^. To assess the role of the host genes DAXX, BRD4, and LIG3 in the production of infectious virus, we performed siRNA-mediated knockdown using siDAXX, siBRD4, and siLIG3 in iSLK.BAC16 cells, followed by lytic reactivation. We collected cell-free KSHV-containing supernatants from siDAXX, siBRD4, siLIG3, and siASN samples and titered them on iSLK cells^81^. We determined the infection levels in iSLK cells by quantification of GFP+ cells via fluorescence microscopy (Figure 4C). Our results show a significant increase in the production of infectious virions, as determined by the elevated percentage of infection after DAXX and BRD4 knockdown, with approximately 350% and 150% increases in infection compared to siASN control samples. In contrast, siLIG3 treated iSLK.BAC16 cells after reactivation produced less KSHV infectious virus than the siASN control cells (Figure 4D). Additionally, we analyzed iSLK cells infected with KSHV isolated from siDAXX, siBRD4, and siLIG3 samples with or without DOX/NaB reactivation by flow cytometry. Viral infection was determined by GFP+ cells (FITC+) quantification (Figure 4E). Knockdown samples without DOX/NaB reactivation did not produce infectious virus as confirmed by the GFP- iSLK cells. Our results revealed an approximately three-fold increase in GFP+ cells after DAXX knockdown and a two-fold increase in GFP+ cells after BRD4 knockdown compared to the siASN control samples. Conversely, we observed a 70% decrease in GFP+ cells after LIG3 knockdown compared to the siASN control samples (Figure 4F). Overall, our results indicate that DAXX and BRD4 inhibition promotes infectious virion production and release. These findings suggest that DAXX and BRD4 may act as host restriction factors that help maintain latency by negatively regulating KSHV virion production. On the other hand, decreased infectious virion production after LIG3 inhibition suggests that LIG3 is a required host factor for the efficient production of infectious KSHV virions during the lytic cycle.

### BRD4 chemical inhibition by JQ1 enhances KSHV lytic gene expression and promotes virus production

BRD4 was selected for further investigation since it is a clinically actionable target. BRD4 inhibitors are commercially available and have been used in cancer research with promising results^82–84^. Utilizing these existing compounds to investigate KSHV virus–host interactions could uncover novel therapeutic strategies for KS, one of the most common cancers in HIV-positive patients. Therefore, we tested JQ1-mediated chemical inhibition of BRD4. JQ1 competitively binds BRD4 bromodomains, inhibiting its binding to chromatin and further functions^85^. We treated iSLK.BAC16 cells with 100nM of JQ1 for 72 hours with or without DOX and NaB for the last 48 hours to induce lytic reactivation. RNA was extracted, and RT-qPCR was performed to determine the expression levels of KSHV lytic genes. Upon reactivation, the immediate early lytic gene ORF45 relative mRNA expression in iSLK.BAC16 cells was significantly increased after JQ1 treatment compared to the DMSO control (Figure 5A). However, early lytic gene ORF59 relative mRNA expression remained constant after JQ1 treatment compared to the DMSO control (Figure 5B). Similarly, the late lytic gene ORF26 was significantly upregulated by 50% in the JQ1-treated cells compared to the DMSO control (Figure 5C). Notably, late lytic gene K8.1 relative mRNA expression was also significantly increased by 1.5-fold in the JQ1-treated cells compared to the DMSO control (Figure 5D).

**Figure 5.**
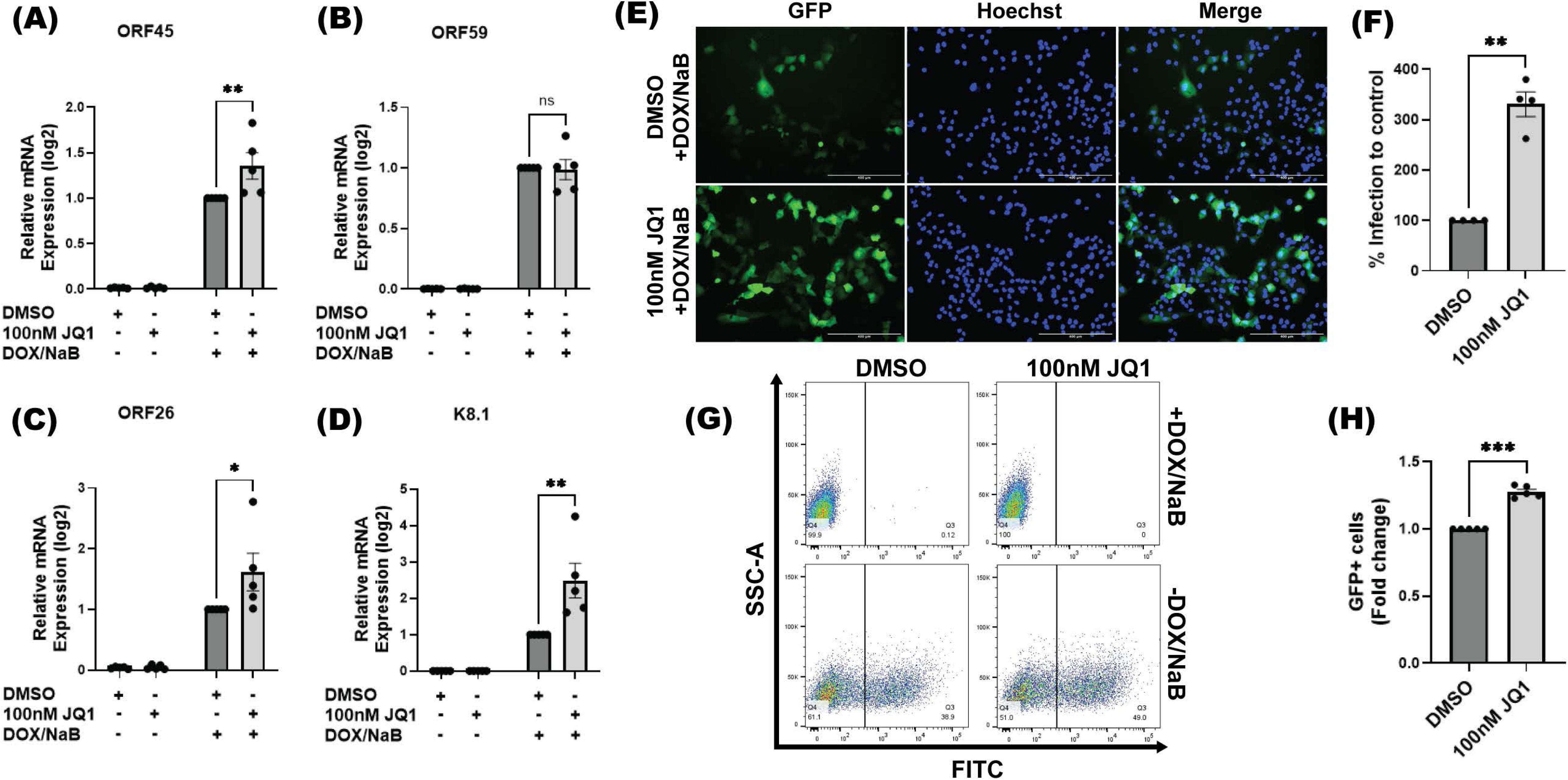
Inhibition of BRD4 by JQ1 promotes increased KSHV viral gene expression and virion production. Relative mRNA expression of the KSHV lytic genes (A) ORF45 (immediate-early lytic), (B) ORF59 (early lytic), (C) ORF26 and (D) K8.1 (late lytic) after 100nM JQ1 treatment in iSLK.BAC16 cells with (+DOX/NaB) or without reactivation (-DOX/NaB). All expression data are relative to Actin (housekeeping) and normalized to DMSO control. n=5. (E) Supernatants of iSLK.BAC16 cells after JQ1 treatment and reactivation were collected, and viral titers were determined in iSLK cells. GFP+ cells correspond to KSHV-infected cells. Titers were assessed at 48 hpi. on an EVOS fluorescent microscope at 10X. (F) Quantification of GFP+ cells observed in (E), n=4. (G) Virus production was also assessed by flow cytometry analysis. Representative dot plot for flow cytometry analysis for GFP+ cells. (H) Quantification of GFP+ cells from (G), n=5. ***, P≤0.001; **, P≤0.01; *, P≤0.05; ns: not statistically significant, Error bars=SEM.

To determine the effect of BRD4 chemical inhibition on the production of KSHV infectious virions, we treated iSLK.BAC16 cells with 100nM JQ1 followed by reactivation. Cell-free KSHV-containing supernatants were collected and titered on iSLK cells. Viral infection was determined by GFP+ cells through fluorescence microscopy, revealing an increase in KSHV infectious virus particles (Figure 5E). Quantification of GFP+ cells confirmed a significant increase in KSHV infectious virus particles after JQ1 treatment in reactivated iSLK.BAC16 cells, with more than 200% increase in infection compared to the DMSO control (Figure 5F). Additionally, we confirmed our viral infection results by flow cytometry analysis. iSLK cells were infected with KSHV isolated from 100nM JQ1-treated and control samples with or without DOX/NaB reactivation (Figure 5G). We determined viral infection by quantifying GFP+ cells (FITC+). Samples without DOX/NaB reactivation did not produce infectious virus (GFP- cells). Our results revealed a 27% increase in GFP+ cells after 100nM JQ1 compared to the control (DMSO) (Figure 5H). Our results showed that chemical inhibition of BRD4 by JQ1 induces upregulation of KSHV lytic gene expression and enhances virus production in iSLK.BAC16 cells upon reactivation. Our findings suggest BRD4 is a potential regulator of KSHV viral latency, controlling the initiation of lytic reactivation.

## DISCUSSION

This study highlights the contribution of DAXX, BRD4, and LIG3 as critical host regulators of KSHV latency and lytic reactivation. By functionally characterizing these LANA-interacting proteins, we provide new insights into the molecular interplay underlying viral persistence and reactivation, with important implications for developing targeted therapies against KSHV-associated diseases. We investigated the role of DAXX, BRD4, and LIG3 in regulating viral gene expression, genome replication, and infectious KSHV virion production using the DOX-inducible iSLK.BAC16 cells, which stably maintain the KSHV genome and support latency and lytic reactivation. Our findings reveal a complex interplay whereby the host factors DAXX and BRD4 act as restriction factors that help maintain viral latency by suppressing lytic reactivation, while the host factor LIG3 functions as a positive regulator that promotes KSHV lytic replication and virus production.

Our IP-MS analysis confirmed the interaction of the host factors DAXX and LIG3 with the KSHV latent protein LANA (Figure 1) in HEK293T cells expressing LANA compared to an empty vector control. These results were consistent with Davis *et al*. (2015)^28^, who mapped the host-KSHV protein interaction network and identified the DAXX-LANA and LIG3-LANA interactions^28^. Interestingly, in our dataset, BRD4 was identified in a LANA-dependent manner in the DAXX immunoprecipitation dataset, raising the possibility that BRD4 detection may result either from its interaction with LANA or from a potential BRD4-DAXX interaction that occurs during the presence of LANA. Despite previous studies have identified the BRD4-LANA interaction^32^, our study reveals for the first time the DAXX-interactome remodeling in the presence of LANA.

Functional depletion of DAXX or BRD4 by siRNA transfection led to a significant upregulation of immediate-early, early, and late lytic gene transcripts upon lytic reactivation in iSLK BAC16 cells, with fold changes ranging from 2.5- to over 37-fold compared to controls (Figure 2). These changes in transcript levels were corroborated by increased viral lytic protein expression by immunofluorescence and western blot analyses (Figure 3), indicating that the knockdowns potentiate lytic reactivation at both the transcriptional and protein levels. Consistent with these results, the knockdown of DAXX or BRD4 resulted in significantly increased intracellular viral DNA copies upon reactivation, indicating that these host factors restrict KSHV lytic genome replication. Importantly, increased production of infectious virions following DAXX and BRD4 knockdown was demonstrated by a significant increase in infection upon titering the cell free supernatant on iSLK cells (Figure 4). Complementing the genetic knockdown data, chemical inhibition of BRD4 with JQ1 recapitulated the enhanced lytic gene expression and virus production phenotype (Figure 5), underscoring BRD4 as a therapeutically targetable regulator of KSHV latency. Fold-changes in gene expression and virus production after JQ1 treatment and lytic reactivation were lower compared to those observed after the knockdown with siBRD4, possibly due to compensatory mechanisms of other bromodomain proteins. Together, our results demonstrated that DAXX and BRD4 act as suppressors of KSHV lytic reactivation, whose inhibition promotes increased viral gene and protein expression, viral replication, and infectious virus production. Interestingly, our IP-MS analysis raises the possibility of a key interaction between DAXX-BRD4, which may contribute to the coordinated regulation of the transition from latency to lytic replication (Figure 6). However, further analyses are required to investigate this potential interaction.

**Figure 6.**
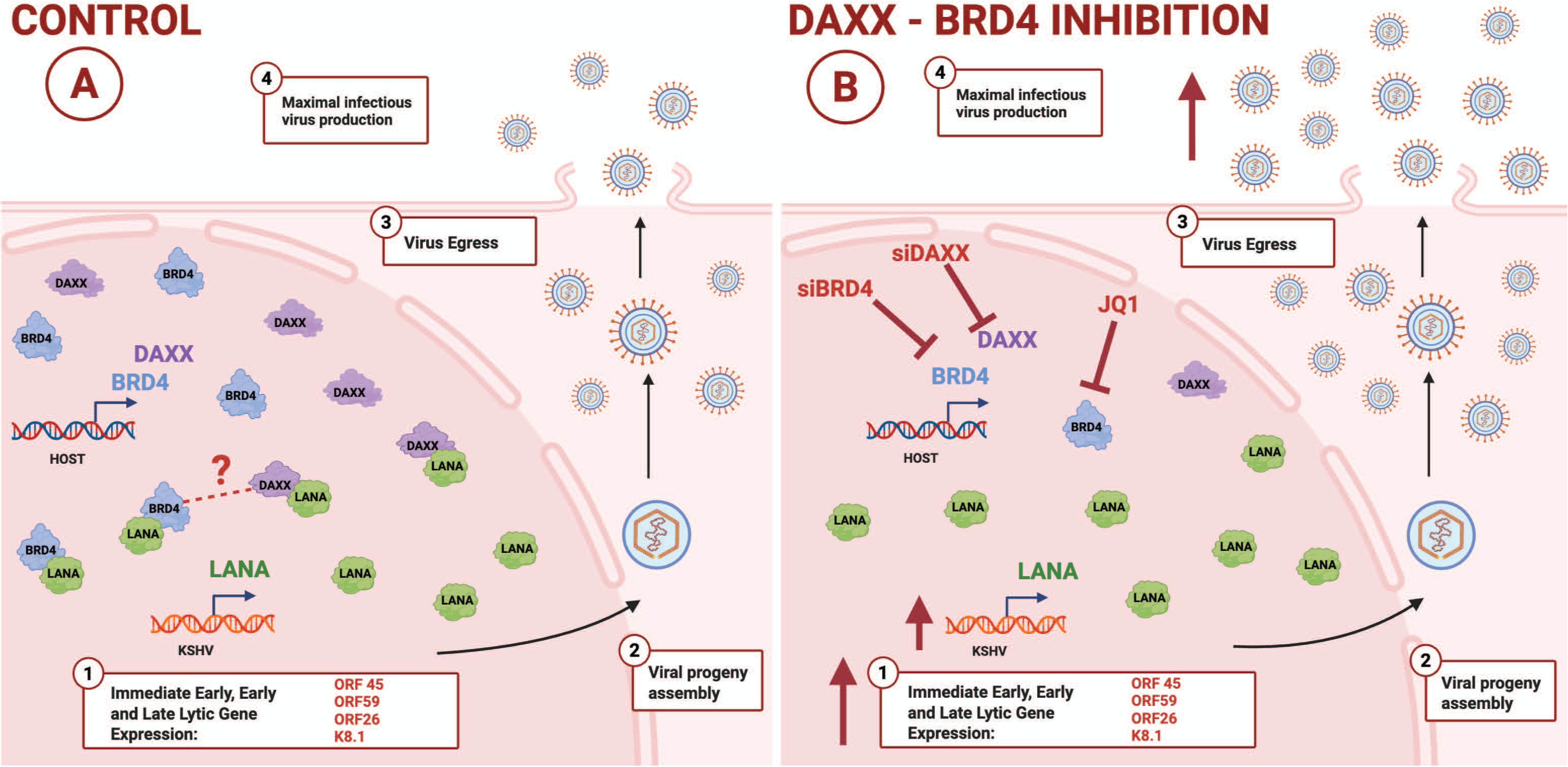
LANA-interacting host proteins DAXX and BRD4 modulate KSHV latency-lytic cycle transition. Inhibition of DAXX and BRD4 in the KSHV reporter cell model iSLK.BAC16 results in increased lytic viral transcripts, proteins, viral replication, and infectious virus production.

In contrast, siRNA-mediated knockdown of LIG3 had the opposite effect. LIG3 depletion caused a significant decrease in KSHV lytic transcript and protein levels (Figure 2**-3)**, diminished viral genome replication, and reduced production of infectious virions. These results implicate LIG3 as a positive effector of KSHV lytic replication, potentially through its known role in DNA repair and genome integrity maintenance, which may facilitate efficient viral DNA replication during lytic reactivation.

Our findings suggest that the LANA-associated host factors examined here play distinct roles in regulating the equilibrium between KSHV latency and lytic reactivation. DAXX and BRD4 seem to act as molecular inhibitors on viral reactivation, possibly by strengthening chromatin-based repression, consistent with their known roles in transcriptional regulation and chromatin organization. Specifically, DAXX has been shown to maintain a repressive chromatin state on viral genomes, as observed in HSV infection. It acts as a histone chaperone for the H3.3 histone, depositing it onto the viral DNA upon initial infection and restricting the ability of the virus to decompact its genome and begin transcription^86^. Interestingly, a previous study showed that DAXX knockdown induced an increased KSHV latent infection, possibly due to the disruption of the host cell nuclear domain 10 (ND10) defense complex, which represses viral gene expression^87^. Additionally, DAXX localizes to LANA nuclear bodies during KSHV latency. DAXX association with LANA nuclear bodies and viral DNA helps maintain viral episomal structure. DAXX is evicted from the LANA bodies during the transition to the lytic cycle^29,49^. Taken together, our results suggest that although DAXX contributes to the maintenance of the episome during latency, its inhibition may facilitate the chromatin decompaction and recruitment of the replication machinery required for efficient lytic reactivation.

In KSHV-infected BCBL1 lymphoma cells, BRD4 has been shown to associate with KSHV episomes at latency control regions, where its interaction with LANA at the terminal repeats (TRs) stabilizes DNA loop structures linking latent and lytic control regions. BRD4 inhibition destabilizes these looped interactions, triggering viral reactivation^88^. This mechanism may explain the enhanced lytic replication observed following BRD4 knockdown in our study.

LIG3 supports host DNA replication^89,90^, and may also assist in the repair and processing of viral DNA intermediates essential for genome replication and virion production. KSHV is known to modulate the host DNA damage response (DDR) by relocalizing or sequestering DNA repair proteins, thereby restricting their ability to activate antiviral inflammatory signals and supporting lytic reactivation^91,92^. In addition, overexpression of the early lytic KSHV protein ORF57 in epithelial cells increased LIG3 protein levels, along with other components of the Non-homologous end joining (NHEJ) repair pathway, suggesting a possible activation of the DNA double-strand break repair during infection^93^. Consistent with this, siRNA-mediated depletion of LIG3 in our experiments resulted in a substantial reduction of viral transcripts and genome replication. Together with the observed decrease in viral proteins and infectious virus production, likely due to the inefficient intracellular viral DNA replication, these findings suggest that KSHV could hijack the host DNA repair machinery, particularly LIG3, to ensure efficient virus production.

From a clinical perspective, the identification of BRD4 as a targetable host factor, whose inhibition by JQ1 promotes lytic reactivation and virion production, opens compelling possibilities for therapeutic intervention. Pharmacological inhibition of BRD4 could be exploited to induce lytic reactivation in KSHV-associated malignancies, thereby sensitizing latently infected cells to antiviral agents or immune clearance. Oncolytic therapeutic approaches for PEL have investigated the use of JQ1 in combination with PEP005 (ingenol-3-angelate), an FDA-approved drug for the topical treatment of actinic keratosis with promising effects on cancer cell growth suppression. This combination treatment induced lytic cycle reactivation in latently infected B-cells and prevented tumor growth^94^. Another study demonstrated that the inhibition of BRD4 by JQ1 in combination with histone deacetylases (HDACs) inhibitors, and the use of small molecule inhibitors targeting both HDAC and BRD4, enhanced viral replication and inhibited the growth of KSHV-positive lymphomas, supporting their potential as therapeutic strategies for PEL^41^. Given our findings that inhibition of BRD4 or DAXX triggers KSHV lytic reactivation, targeting either host factor individually or in combination may provide a broader therapeutic platform that could be applied to other KSHV-associated diseases.

Our results from gene and protein expression analyses, viral genome quantification, and infectious virus production collectively support a model in which DAXX and BRD4 function antagonistically to LIG3 in regulating KSHV replication cycles. Further investigation into the molecular mechanisms governing these interactions, including chromatin remodeling, transcriptional complex dynamics, and DNA repair coordination, will deepen our understanding and may facilitate the design of novel antiviral strategies for KSHV-associated diseases. Altogether, this study advances the functional framework that links host factors to the regulation of KSHV latency and reactivation.

## Acknowledgements

We would like to thank all the members of the Sanchez lab for their creative input throughout this study. We would further like to acknowledge and thank the Biology Department at the University of Texas at Dallas for helpful discussion on this project. We thank Dr. Nevan Krogan and additional members of the Krogan group for helpful comments and discussion.

## Notes

### Competing Interest Statement

The authors have declared no competing interest.

